# A 2,000-year-old specimen with intraerythrocytic *Bartonella quintana*

**DOI:** 10.1101/2020.03.13.990580

**Authors:** R Barbieri, B-H-A Mai, T Chenal, M-L Bassi, D Gandia, L Camoin-Jau, H Lepidi, G Aboudharam, M Drancourt

## Abstract

Photogrammetry and cascading microscopy investigations of dental pulp specimens collected from 2,000-year-old individuals buried in a Roman necropolis in Besançon, France, revealed unprecedented preserved tissular and cellular morphology. Photogrammetry yielded 3-D images of the smallest archaeological human remain ever recovered. Optical microscopy examinations after standard hematoxylin-phloxine-saffron staining and anti-glycophorin A immunohistochemistry exposed dental pulp cells, in addition erythrocytes were visualized by electron microscopy, which indicated that the ancient dental pulp trapped a blood drop. Fluorescence *in situ* hybridization applied on red blood cells revealed the louse-borne pathogen *Bartonella quintana*, a finding confirmed by polymerase chain reaction assays. Through paleohistology and paleocytology, we demonstrate that ancient dental pulp preserved intact blood cells at the time of the individual’s death, offering an unprecedented opportunity to engage in direct and indirect tests to diagnose pathogens in ancient buried individuals.

## INTRODUCTION

Teeth are organs recoverable in human individuals as old as ~ 300,000 years ^1^ and could be the last organs that remain intact from dead individuals. Teeth contain dental pulp on which fruitful paleomicrobiology investigations have been performed ^2^. Indeed, ancient dental pulp preserves ancient pathogen biomolecules, including mycolic acids ^3^ and DNA, under favorable biochemical conditions ^4^. From the ancient dental pulp, genome-wide data were analyzed from various micro-organisms including bacteria, viruses and parasites, such as the plague agent *Yersinia pestis* ^5–17^, the relapsing fever agent *Borrelia recurrentis* ^18^, the leprosy agent *Mycobacterium leprae* ^19–21^, the typhoid fever pathogen *Salmonella enterica* ^22,23^, the hepatitis B virus ^24,25^ and the malaria agent *Plasmodium falciparum* ^26^. In addition, dental pulp preserves ancient proteins in such a way that the paleoproteomics of pathogens can be performed ^27^. Recently, investigations unexpectedly revealed host peptides, including peptides derived from conjunctive dental pulp tissue and plasmatic peptides, such as coagulation factors and immunoglobulins ^27^. These observations, validated by the establishment of paleoserology protocols for the detection of immunoreactive immunoglobulins (Drancourt M., unpublished data), indicated that ancient dental pulp can preserve blood and its serum phase.

Here, using paleocytology, we show that ancient dental pulp also can preserve blood cells exhibiting unanticipated and perfectly conserved morphology. Illustrating one outcome of this discovery in the field of paleomicrobiology, we performed microscopic-only detection of one ancient intracellular pathogen, *Bartonella quintana* ^28^, in a 2,000-year-old Roman dental pulp specimen obtained in France.

## RESULTS

### Photogrammetry

A total of 25 teeth were collected from five individuals (here referred to as individuals 17, 20, 21, 33 and 35) discovered in a 2,000-year-old Roman archaeological site (Viotte Nord, Besançon) in France (Supplementary Information). The dental pulp was recovered from each tooth as previously described, and two dental pulp aliquots were prepared ^29,30^ (Supplementary Video 1). The dental pulp recovered from individual 17, being macroscopically intact, was further analyzed by photogrammetry. A total of 1,500 images enabled us to recreate the pulp sample in full volume at the following photogrammetry specifications: a scattered cloud of 104.256 points, a dense cloud of 361.35 points, a mesh volume of 822.771 faces, and finally a mosaic texturing of 12.288 × 12.288. The quality of each photo thus made it possible to observe the dental pulp, with a rendering that was close to being observable with the naked eye or binocular glasses, as shown in Figure 1 (Supplementary Video 2).

**Figure 1.**
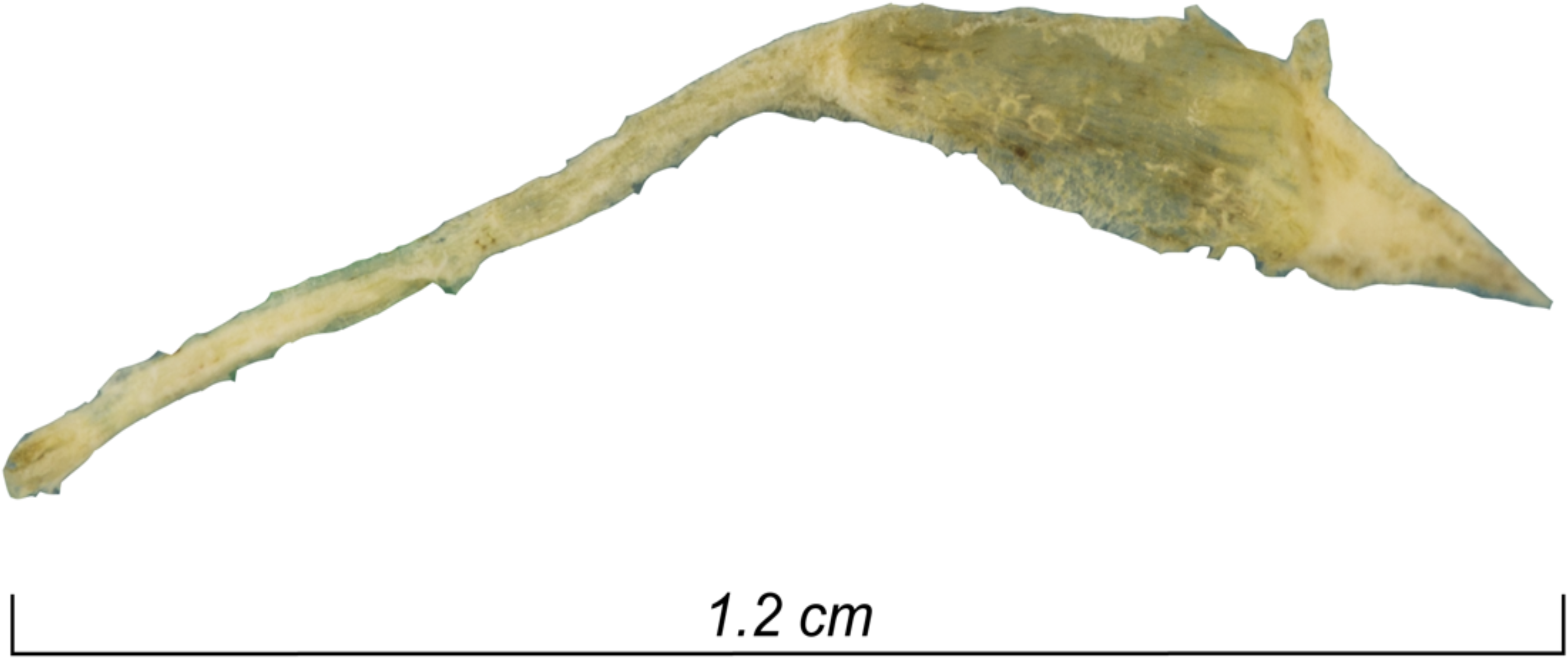
Photogrammetry of a 2,000-year-old intact dental pulp recovered from Ind. 17.

### qPCR-based detection of *B. quintana*

A total of 24 dental pulp specimens collected from individuals 20, 21, 33 and 35 were tested for the presence of *B. quintana* DNA by detecting the *yop*P ^31^ and intergenic transcribed spacer (ITS) ^32^ sequences by real-time PCR, as previously described ^30^. With the negative controls remaining negative, one dental pulp sample collected from individual 35 was positive for two of the molecular targets, 2/24 dental pulp samples were positive for *yop*P only, and 7/24 dental pulp samples were positive for the ITS region only. Therefore, individual 35 was determined to be *B. quintana*-positive; individual 33 was determined to be *B. quintana*-negative (Table 1); and the pulp from both individuals was further investigated by microscopy as described.

**Table 1.** Overview of *yopP* and ITS detection in samples from individuals 20, 21 and 35. Dental pulp specimens positive for *yopP* or/and ITS are presented in green while negative specimens are presented in red.

| Indivudual | N° Tooth | *yopP* | ITS | Cq |
| --- | --- | --- | --- | --- |
| 21 | 1 |  |  | 37.8 |
|  | 2 |  |  | 37.4 |
|  | 3 |  |  | 38.2 |
|  | 4 |  |  | 33.2 |
|  | 5 |  |  | 34.2 |
| 20 | 8 |  |  | 35.3 |
|  | 9 |  |  | 34.4 |
|  | 10 |  |  | 34.8 |
| 35 | 13 |  |  | 36.2 |
|  | 14 |  |  | 33.4 ; 34.5 |

### Paleocytology

The dental pulp specimen collected from the *B. quintana*-positive individual (35) was immersed in a rehydration buffer, adapted from Sandison el al.^33^, consisting of 1% formalin, 96% ethanol and 5% ammonium bicarbonate for 24 hours before hematoxylin-phloxine-saffron (HPS) staining and anti-glycophorin-A staining, which is specific for detecting erythrocytes ^34,35^. The microscopic observation was performed from 10× to 100× magnification. In these experiments, the dental pulp sample collected from the *B. quintana*-negative individual (33 was used as the negative control and manipulated strictly in parallel to the one the dental pulp sample collected from the *B. quintana*-positive individual (35). Microscopic observations of the HPS-stained dental pulp specimens (5 slides for individual 35 and 5 slides for individual 33) unexpectedly yielded the presence of pink or red cells with sizes and morphological characteristics consistent with those of erythrocytes (Fig. 2A). Anti-glycophorin A staining (10 slides for individual 35 and 10 slides for individual 33) (Fig. 2B) yielded red-orange-brown colored cells devoid of nuclei, ranging in size from 2 to 8 μm, confirming these cells as erythrocytes. The stained erythrocytes presented different shapes, ranging from round to square, and some cells presents concave surfaces specific of erythrocytes, while these morphological characteristics were not observed in the negative controls obtained through the use of an irrelevant antibody. Ancient erythrocytes appeared as separate cells or agglutinated in erythrocytic islets (Fig. 2). Electron microscopy confirmed the remarkable conservation of the erythrocyte morphology 2,000 years after the death of the individual (Fig. 3B).

**Figure 2.**
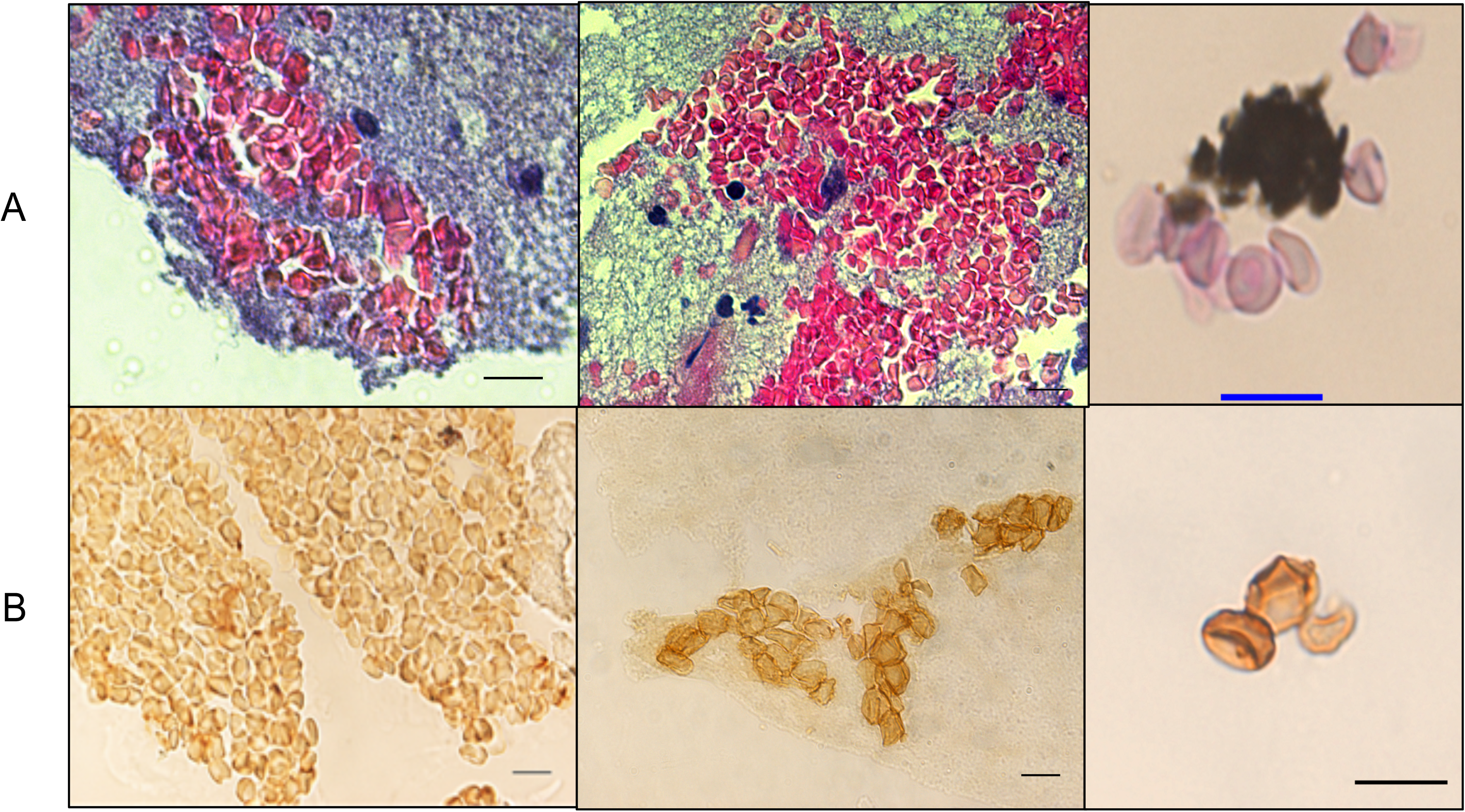
Identification of ancient erythrocytes in individual 33 (*B. quintana*-negative) and individual 35 (*B. quintana*-positive) A. HPS staining (10-μm scale) and B. anti-glycophorin A staining (10-μm scale).

**Figure 3.**
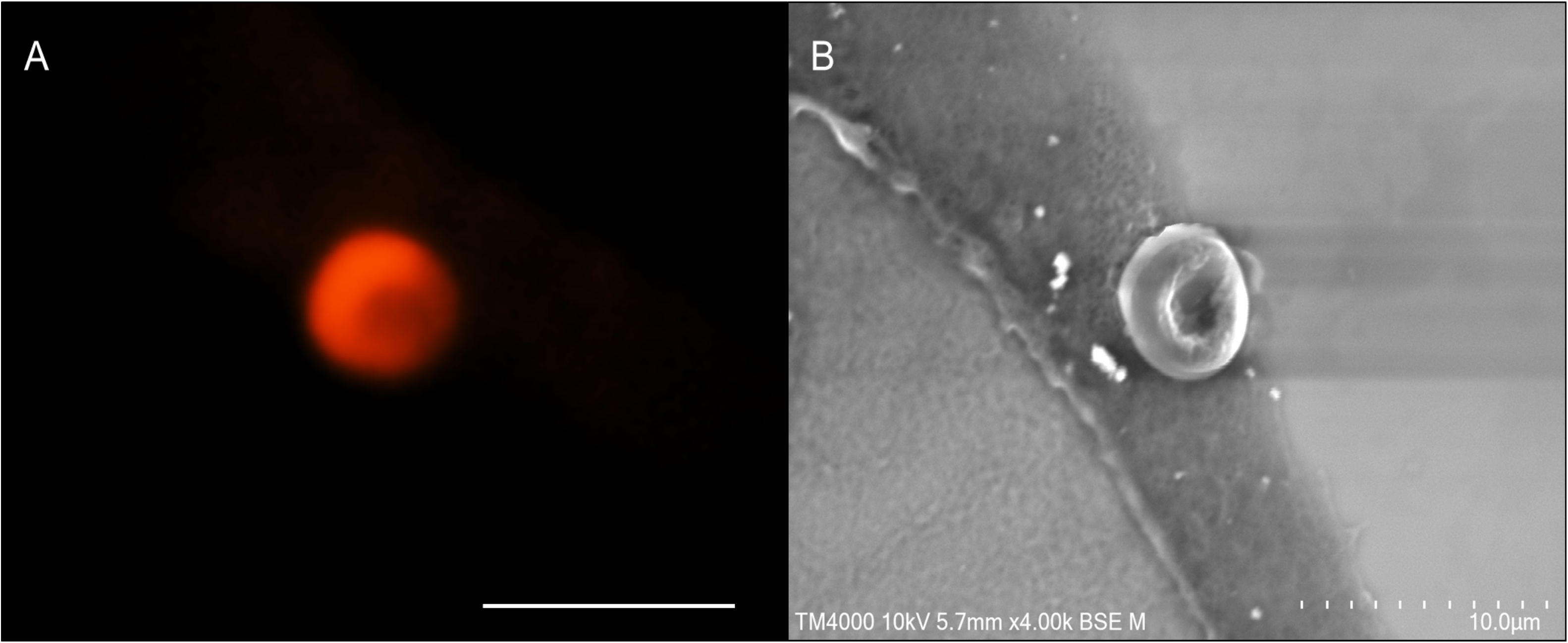
A. Fluorescence microscopy of an erythrocyte in Ind. 35 using wavelength 555 nm. B. The same erythrocyte was observed by scanning electron microscopy (10-μm scale).

### Fluorescence *in situ* hybridization of *B. quintana* in ancient erythrocytes

The detection of morphologically intact erythrocytes in the dental pulp of two individuals, 33 (negative control) and 35 (*B. quintana* positive), provided a unique opportunity to test for microscopically identifiable intraerythrocytic *B. quintana* organisms using fluorescence *in situ* hybridization (FISH). Accordingly, the dental pulp samples were stained using 4’,6’-diamidino-2-phenylindole (DAPI), a DNA-binding dye, here used to screen erythrocytes for the presence of exogenous, presumably bacterial, DNA. Indeed, *B. quintana* is acknowledged as such an intraerythrocytic organism ^36^, which has been previously detected in ancient dental pulps and bones collected in several archaeological sites in France ^37–41^. Erythrocyte autofluorescence (anticipated to interfere with the FISH detection of *B. quintana*) was not observed in erythrocytes, which exhibited a flat morphology (Fig. 4). Confocal microscopy revealed that one such flat erythrocyte in individual 35 had been stained blue with DAPI, stained red with the pan-bacterial probe and stained green with the *B. quintana*-specific probe, remaining dark with the non—specific probe, whereas no such images were observed in the sample from negative-control individual 33, indicating that bacteria were present in the erythrocytes of individual 35 and that these bacteria were *B. quintana* (Fig. 4).

**Figure 4.**
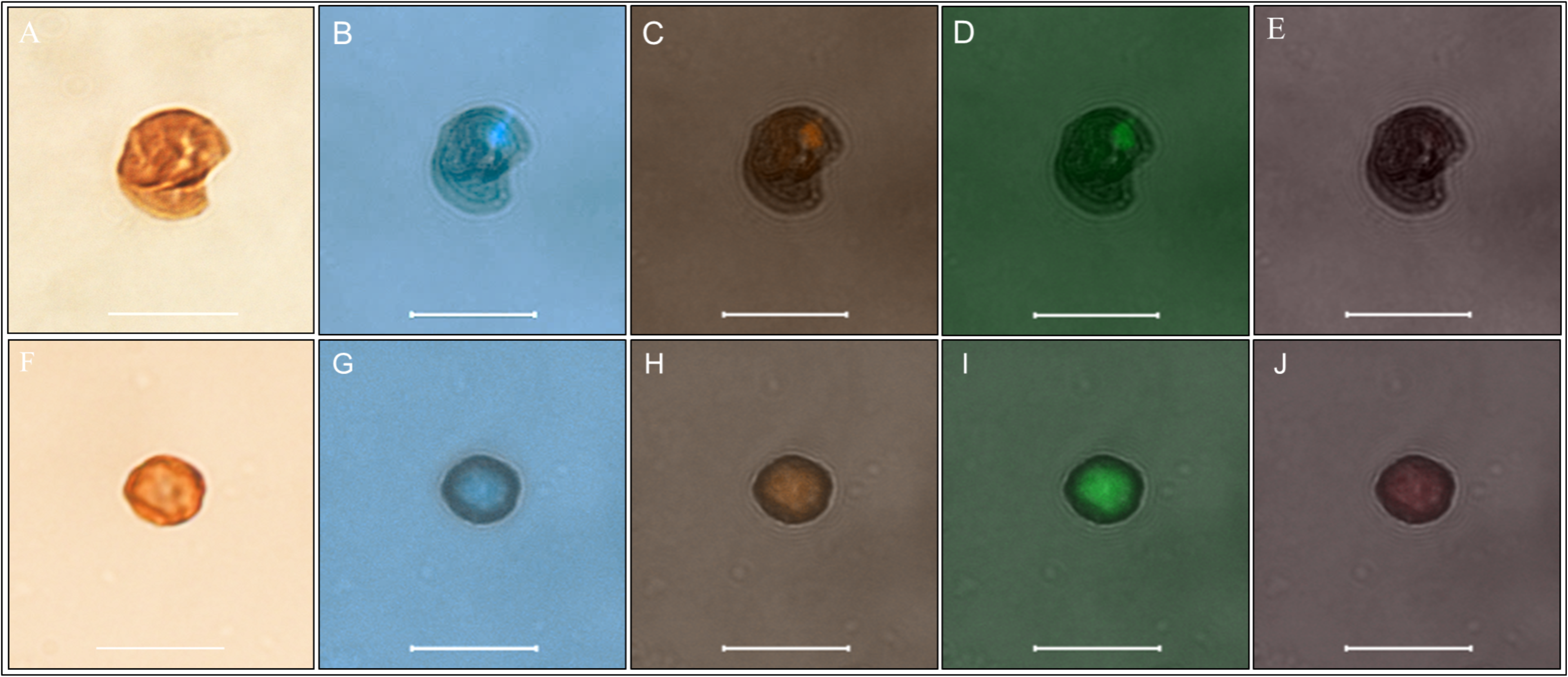
FISH revealed *B. quintana*-infected erythrocytes from individual 35, with a *B. quintana*-negative autofluorescent erythrocyte from individual 33 used as a negative control. A and F: Optical microscopy; B and G: confocal microscopy with DAPI staining. C and H: confocal microscopy with EUB probe. D and I: Confocal microscopy with a *B. quintana*-specific probe. E and J: Confocal microscopy with nonEUB probe.

**Figure 5.**
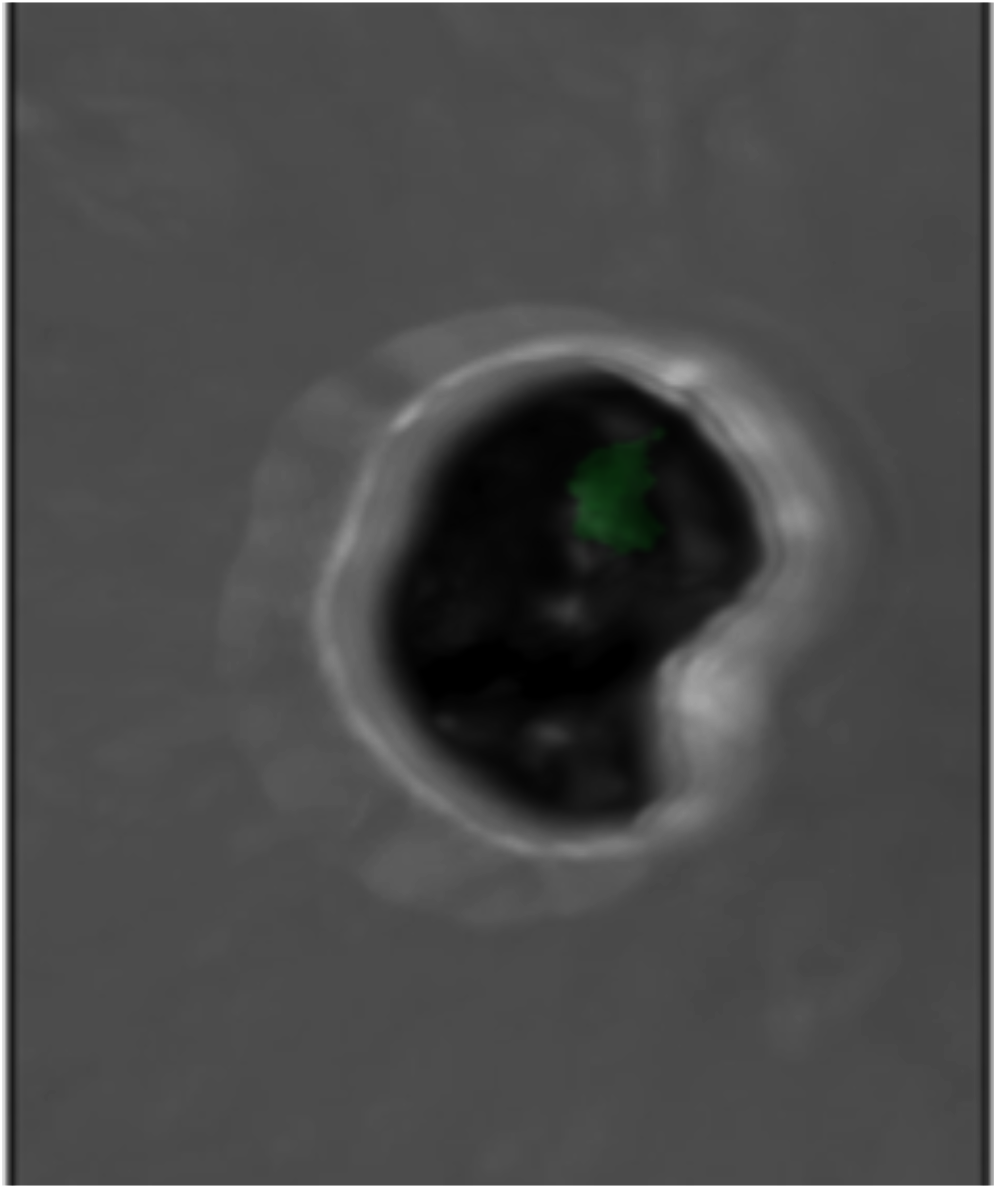
3D-FISH revealed *B. quintana* inside individual 35 erythrocytes under a green channel-specific probe.

## DISCUSSION

Microscopy has been the first ever laboratory tool used by the 19th century for the diagnosis of infectious (bacterial) diseases by the direct observation of microbes in clinical samples ^42,43^, and we are now reporting that microscopy is still a diagnostic approach of value when addressing pathogens in ancient specimens. In this study, all microscopic observations were authenticated by negative controls. Additionally, microscopic detection of *B. quintana* was confirmed by two different qPCR assays, including negative controls and no positive controls.

In this study, the intact preservation of dental pulp blood cells is consistent with few previous observations. Indeed, human erythrocytes have been detected from mummified soft tissues as old as 5,300 years old ^33,44–47^, in one 1,600-year-old bone marrow ^48^ and in Medieval bone remains ^49^. Erythrocytes of unknown source have also been microscopically detected on prehistoric archeological tools potentially used for hunting and meat preparation ^50,51^.

Paleocytological observations reported here add one more specimen (the dental pulp) in which to base the microscopic detection of mammal cells, here human erythrocytes, further illustrating that archeological specimens could preserve morphological intact blood cells, in addition to serum proteins ^52^. Here, we show that ancient teeth entrapped a blood clot following the death of the individual. Combining these observations suggested that indirect (serological) and direct diagnosis tests using blood could be assayed on ancient dental pulps.

As an illustration, paleocytology unanticipatedly allowed microscopic detection of a 2,000 year-old intraerythrocytic *B. quintana* in the same way that it has been previously reported in modern blood smears ^36^. Our study therefore adds a new approach for the detection of this pathogen that, so far, has been detected only based on PCR-based techniques (Table 1). The efficiency of PCR-based approaches could be limited by a risk of laboratory cross-contamination leading to false positives results and the presence of PCR inhibitors leading to false negatives results ^53^. Microscopic techniques including FISH are less prone to such limitations, offering a complementary approach to PCR-based techniques. The observations reported here pave the way to sorting *B. quintana*-infected red blood cells for further characterization of ancient pathogens, including whole genome sequencing.

This report introduces paleocytology as an innovative approach for the analysis of ancient dental pulp to study both host- and host-associated pathogens, complementing the study of ancient biomolecules ^2,3,27,54^.

## MATERIALS AND METHODS

### The archaeological site

The preventive excavation “Viotte Nord” was carried out as part of the work related to a railway station in Besançon in northeastern France. The archaeological intervention took place on a surface of 2,786 m2 on which forty burials, six secondary cremation burials and a probable dump area were exhumed. The furniture from funeral structures, as well as C^14^ dating on bones and charcoal, in this burial site was attributed to the transition period between the High Empire and the Lower Empire, i.e., I-IV centuries AD (Supplementary Figure 1). The excavation of funerary materials revealed 43 more-or-less well-preserved skeletons. Most individuals were wrapped in textiles in a wooden container; some individuals were accompanied by grave goods, including ceramics, glass, shoes and animal bones. We received teeth from four individuals collected from three burials (SEP): SEP 230 (Supplementary figure 2) was a double-burial site containing one mature man (Ind. 21) and one woman (Ind. 20), with archaeological studies and C^14^ dating, this structure was dated between the 2^nd^ and 3^rd^ centuries; SEP 283 (Supplementary figure 3) contained an immature individual (Ind. 33), with archaeological studies dating the structure between the middle 2^nd^ century and the 4^th^ century; SEP 286 (Supplementary figure 4) contained a young adult (Ind. 35) of undetermined sex, archaeological studies dating the structure to the 2^nd^ century; SEP 225 (Supplementary figure 5) contained a mature adult man (Ind. 17) of approximately 1.74-m stature, C14 analysis dating this individual from between the second and third centuries (Fig. 2 and Supplementary information). We analyzed a total of 25 teeth, including seven teeth collected from Ind. 21, five teeth collected from Ind. 20, eight teeth collected from Ind. 35, four teeth collected from Ind. 33 and one tooth collected from Ind. 17.

### Pulp collection and photogrammetry

Teeth collected from 5 individuals at the site of Viottes were then processed in a dedicated laboratory room at the Institute of IHU Méditerranée Infection, Marseille, France. The teeth were brushed under tap water to remove dirt and debris and finally washed with nuclease-free water. Under a chemical hood, an electric motor with a rotating diamond disc was used to open each tooth longitudinally, dividing it into two parts. Pulp was extracted using a sterile dentist excavator and scraped into two sterile Eppendorf tubes: one was maintained for use DNA extraction as previously described ^29,30^ (Supplementary video 2) and the other was used to prepare microscopic slides as described below. The dental pulp from individual 17 was preserved in such good condition that it was analyzed by photogrammetry. To digitize the image as close as possible to the true ancient dental pulp, reputed to be transparent, we used precision photogrammetry material (Agisoft Photoscan 1.5, Agisoft LLC, Saint Petersburg, Russia) using a 100-mm macro lens and a full frame sensor (EOS 6D Mark II, Canon, Tokyo, Japan). Because of the transparency and smallness of the object, a white light room was installed in the laboratory to obtain controlled light diffusion, as well as to capture images in three focused sectors. A single in-focus snapshot of the entire object using a bracketing technique was generated from a combination of three snapshots developed based on three separate parts of the object.

### DNA extraction and amplification

DNA was extracted with a modified phenol-chloroform protocol ^55^. Briefly, each Eppendorf tube containing 5-15 mg dental pulp powder was added to a solution of 10 μL of 25 mg/mL proteinase K, 10 μL of 10% dodecyl sulfate sodium (SDS), and 200 μL of sterile water and agitated at 56°C overnight before the addition of 220 μL of stabilized phenol/chloroform/isoamyl-alcohol (25:24:1) (Biosolve, Valkenswaard, Netherlands). The tube was centrifuged at 16,000 g for 5 min, and 220 μL of phenol/chloroform/isoamyl-alcohol was mixed with the upper phase for a second extraction. The upper phase was incubated at −20°C overnight with a solution of 1 μL of glycogen, 44 μL of ammonium acetate (10 M) and 440 μL of absolute ethanol. After a 30-min centrifugation at 20,000 g at room temperature, the pellet was washed with 440 μL of 70% ethanol%, dried at 50°C for 10 min and finally resuspended in 50 μL of nuclease-free water. the DNA extracts were tested for *B. quintana* DNA using quantitative real-time PCR (qPCR) targeting the ITS region and *yop*P genes sequences, as previously described ^56^. A dental pulp sample was considered positive when the qPCR reaction was positive with a cycle number (Ct) lower than 40 for at least one gene. The qPCR results guided the choice of one *B. quintana*-positive dental pulp and one *B. quintana*-negative dental pulp sample for subsequent analyses (Table 1).

### Rehydration and slide preparation

The rehydration process was adapted from the protocol originally reported by Sandison [Sandison 1955]. Briefly, 200 μL of a solution containing the dental pulp and 2 volumes 5% ammonium bicarbonate, 5 volumes of 1% formaldehyde and 3 volumes 96% ethanol was maintained for 24 hours at room temperature. Then, the supernatant was removed following centrifugation at 16.000 g for 5 min on a bench-top Eppendorf centrifuge. Dental pulp pellets were collected, fixed in 10% buffered formalin, dehydrated in grade alcohol and embedded in paraffin. Serial 3.5-μm sections were cut from the block and mounted on poly-L-lysine-coated glass slides.

### Anti-glycophorin A and HPS staining

To detect erythrocytes, the paraffin sections were incubated with anti-glycophorin A antibody JC 159 (Mouse Monoclonal Antibody, ref: Mob 066-05, Diagnostic BioSystems, Nanterre, France) at a 1/500 dilution using a Ventana Benchmark autostainer (Ventana Medical Systems, Inc., Tucson, AZ). Briefly, after removing the paraffin and washing them with reaction buffer, the sections were incubated with a primary antibody for 32 min at room temperature and then incubated with the reagent from an iVIEW DAB detection kit (Roche Diagnostics, Meylan, France). The sections were counterstained with hematoxylin and post-counterstained with bluing reagent. A negative control incorporating an irrelevant monoclonal antibody was run in parallel. The other paraffin sections were stained with HPS on a Tissue-Tek Prisma autostainer (Sakura, CA, US) and microscopically examined (DM2500 microscope Leica, Vienna, Austria).

### Fluorescence microscopy and transmission electron microscopy

Paraffin was removed from the slices by incubating them for 15 min at 65°C followed by 10 min at room temperature in a xylene substitute solution (SafeSolv, Fontenay-sous-Bois, France). The tissue section was emerged in a descending ethanol series (100%, 90%, and 70%; 5 min each), rinsed in sterile water and air-dried. The slides were covered with cover slips, sealed with flexible mounting adhesive (Fixogum, Marabu, Germany) and examined with a confocal microscope (LSM800 microscope, Zeiss, Marly-le-Roi, France). After the coverslips were removed, the tissues were observed by scanning microscopy (TM4000 Plus microscope, Hitachi, Tokyo, Japan).

### Fluorescence *in situ* hybridization (FISH)

We used four probes as previously described ^57^: a 16S rRNA gene sequence-based probe (5’-AATCTTTCTCCCAGAGGG) specific for *B. quintana*, which was labeled with Alexa-488 (Eurogentec, Angers, France); an Alexa-555-labeled probe (Eurogentec); a universal bacterial 16S rRNA gene sequence-based probe EUB338 (5′-GCTGCCTCCCGTAGGAGT), used as a positive control; and an Alexa-647-labeled (Eurogentec) nonspecific probe non-EUB (5′-ACTCCTACGGGAGGCAGC), used as a negative control. The same slides used for anti-glycophorin A immunohistochemistry were used to perform FISH. The slides were covered with 10 μL of a solution containing 1 μL of the 16S probe (10 μmol/L), 1 μL of the EUB-338 probe (10 μmol/L), 1 μL of the non-EUB probe (10 μmol/L) and 1 μL of a solution containing 0.1% Tween 20, 5 μL of hybridization buffer (0.9 M NaCl; 20 mM Tris-HCL, pH 8.0; 30% formamide; and 0.01% SDS) and 1 μL of distilled water. The slides were covered with a coverslip, sealed with adhesive and incubated for 10 min at 65°C and overnight at 37°C on a FISH-hybridizer (Dako; Agilent Technologies, Santa Clara, CA). The slides were then immersed in a series of baths with SSC buffer at different concentrations of 4×, 2×, 1×, and 0.5×, for 5 min in each bath, and finally, rapidly immersed in a distilled water bath at room temperature. The air-dried slides were stained with DAPI (ProLong Diamond antifade - Fisher Scientific) and examined with a Zeiss LSM800 confocal microscope. The fluorescence of the *B. quintana*-specific probe was read with a green filter, the universal probe EUB338 with an orange filter, the non-EUB probe with a red filter and DAPI with a blue filter. Bright-field images were obtained with an ESID detector excited with an FITC 488-nm laser, images in white and black were generated by using combinations of color channels. Single images were generated from the acquired 4-μm-thick Z-stack images.

## REFERENCES

1. Hublin, J.-J. et al. New fossils from Jebel Irhoud, Morocco and the pan-African origin of Homo sapiens. Nature 546, 289–292 (2017).

2. Drancourt, M. & Raoult, D. Palaeomicrobiology: current issues and perspectives. Nature Reviews Microbiology 3, 23–35 (2005).

3. Gernaey, A. M. et al. Mycolic acids and ancient DNA confirm an osteological diagnosis of tuberculosis. Tuberculosis 81, 259–265 (2001).

4. Turner-Walker, G. The chemical and microbial degradation of bones and teeth. Advances in human palaeopathology 592, (2008).

5. Spyrou, M. A. et al. Analysis of 3800-year-old Yersinia pestis genomes suggests Bronze Age origin for bubonic plague. Nat Commun 9, 2234 (2018).

6. Spyrou, M. A. et al. Historical Y. pestis Genomes Reveal the European Black Death as the Source of Ancient and Modern Plague Pandemics. Cell Host Microbe 19, 874–881 (2016).

7. Spyrou, M. A. et al. Phylogeography of the second plague pandemic revealed through analysis of historical Yersinia pestis genomes. Nature Communications 10, 1–13 (2019).

8. Damgaard, P. de B. et al. 137 ancient human genomes from across the Eurasian steppes. Nature 557, 369–374 (2018).

9. Rascovan, N. et al. Emergence and Spread of Basal Lineages of Yersinia pestis during the Neolithic Decline. Cell 176, 295–305.e10 (2019).

10. Rasmussen, S. et al. Early divergent strains of Yersinia pestis in Eurasia 5,000 years ago. Cell 163, 571–582 (2015).

11. Andrades Valtueña, A. et al. The Stone Age Plague and Its Persistence in Eurasia. Curr. Biol. 27, 3683–3691.e8 (2017).

12. Bos, K. I. et al. A draft genome of Yersinia pestis from victims of the Black Death. Nature 478, 506–510 (2011).

13. Bos, K. I. et al. Eighteenth century Yersinia pestis genomes reveal the long-term persistence of an historical plague focus. Elife 5, e12994 (2016).

14. Namouchi, A. et al. Integrative approach using Yersinia pestis genomes to revisit the historical landscape of plague during the Medieval Period. Proc. Natl. Acad. Sci. U.S.A. 115, E11790–E11797 (2018).

15. Keller, M. et al. Ancient Yersinia pestis genomes from across Western Europe reveal early diversification during the First Pandemic (541-750). Proc. Natl. Acad. Sci. U.S.A. 116, 12363–12372 (2019).

16. Wagner, D. M. et al. Yersinia pestis and the plague of Justinian 541-543 AD: a genomic analysis. Lancet Infect Dis 14, 319–326 (2014).

17. Feldman, M. et al. A High-Coverage Yersinia pestis Genome from a Sixth-Century Justinianic Plague Victim. Mol. Biol. Evol. 33, 2911–2923 (2016).

18. Guellil, M. et al. Genomic blueprint of a relapsing fever pathogen in 15th century Scandinavia. Proceedings of the National Academy of Sciences 115, 10422–10427 (2018).

19. Schuenemann, V. J. et al. Genome-wide comparison of medieval and modern Mycobacterium leprae. Science 341, 179–183 (2013).

20. Schuenemann, V. J. et al. Ancient genomes reveal a high diversity of Mycobacterium leprae in medieval Europe. PLoS pathogens 14, (2018).

21. Krause-Kyora, B. et al. Ancient DNA study reveals HLA susceptibility locus for leprosy in medieval Europeans. Nature communications 9, 1–11 (2018).

22. Vågene, Å. J. et al. Salmonella enterica genomes from victims of a major sixteenth-century epidemic in Mexico. Nature ecology & evolution 2, 520–528 (2018).

23. Zhou, Z. et al. Pan-genome analysis of ancient and modern Salmonella enterica demonstrates genomic stability of the invasive para C lineage for millennia. Current Biology 28, 2420–2428. e10 (2018).

24. Krause-Kyora, B. et al. Neolithic and medieval virus genomes reveal complex evolution of hepatitis B. Elife 7, e36666 (2018).

25. Mühlemann, B. et al. Ancient hepatitis B viruses from the Bronze Age to the Medieval period. Nature 557, 418 (2018).

26. Marciniak, S. et al. Plasmodium falciparum malaria in 1st–2nd century CE southern Italy. Current Biology 26, R1220–R1222 (2016).

27. Barbieri, R. et al. Paleoproteomics of the Dental Pulp: The plague paradigm. PLoS ONE 12, e0180552 (2017).

28. Maurin, M. & Raoult, D. Bartonella (Rochalimaea) quintana infections. Clinical microbiology reviews 9, 273–292 (1996).

29. Drancourt, M., Aboudharam, G., Signoli, M., Dutour, O. & Raoult, D. Detection of 400-year-old Yersinia pestis DNA in human dental pulp: an approach to the diagnosis of ancient septicemia. Proc. Natl. Acad. Sci. U.S.A. 95, 12637–12640 (1998).

30. Raoult, D. et al. Molecular identification by ‘suicide PCR’ of Yersinia pestis as the agent of medieval black death. Proc. Natl. Acad. Sci. U.S.A. 97, 12800–12803 (2000).

31. Angelakis, E., Rolain, J.-M., Raoult, D. & Brouqui, P. Bartonella quintana in head louse nits. FEMS Immunology & Medical Microbiology 62, 244–246 (2011).

32. O’Rourke, L. G. et al. Bartonella quintana in cynomolgus monkey (Macaca fascicularis). Emerging infectious diseases 11, 1931 (2005).

33. Sandison, A. T. The histological examination of mummified material. Stain technology 30, 277–283 (1955).

34. Marchesi, V. T., Tillack, T. W., Jackson, R. L., Segrest, J. P. & Scott, R. E. Chemical characterization and surface orientation of the major glycoprotein of the human erythrocyte membrane. Proceedings of the National Academy of Sciences 69, 1445–1449 (1972).

35. Furthmayr, H. Structural comparison of glycophorins and immunochemical analysis of genetic variants. Nature 271, 519–524 (1978).

36. Rolain, J.-M. et al. Bartonella quintana in human erythrocytes. The Lancet 360, 226–228 (2002).

37. Tran, T.-N.-N. et al. High throughput, multiplexed pathogen detection authenticates plague waves in medieval Venice, Italy. PLoS ONE 6, e16735 (2011).

38. Tran, T.-N.-N., Forestier, C. L., Drancourt, M., Raoult, D. & Aboudharam, G. Brief communication: co-detection of Bartonella quintana and Yersinia pestis in an 11th-15th burial site in Bondy, France. Am. J. Phys. Anthropol. 145, 489–494 (2011).

39. Raoult, D. et al. Evidence for louse-transmitted diseases in soldiers of Napoleon’s Grand Army in Vilnius. The Journal of infectious diseases 193, 112–120 (2006).

40. Drancourt, M., Tran-Hung, L., Courtin, J., Lumley, H. de & Raoult, D. Bartonella quintana in a 4000-year-old human tooth. The Journal of infectious diseases 191, 607–611 (2005).

41. Grumbkow, P. v et al. Brief communication: Evidence of Bartonella quintana infections in skeletons of a historical mass grave in Kassel, Germany. American journal of physical anthropology 146, 134–137 (2011).

42. Koch, R. The etiology of tuberculosis. Reviews of infectious diseases 4, 1270–1274 (1982).

43. Pasteur, L. De l’extension de la théorie des germes à l’étiologie de quelques maladies communes. CR Acad Sci 90, 1033–1044 (1880).

44. Zimmerman, M. R. Blood cells preserved in a mummy 2000 years old. Science 180, 303–304 (1973).

45. Riddle, J. M., Ho, K.-L., Chason, J. L. & Schwyn, R. C. Peripheral blood elements found in an Egyptian mummy: a three-dimensional view. Science 192, 374–375 (1976).

46. Janko, M., Stark, R. W. & Zink, A. Preservation of 5300 year old red blood cells in the Iceman. Journal of the Royal Society Interface 9, 2581–2590 (2012).

47. Reyman, T. A., Barraco, R. A. & Cockburn, A. Histopathological examination of an Egyptian mummy. Bulletin of the New York Academy of Medicine 52, 506 (1976).

48. Setzer, T. J., Sundell, I. B., Dibbley, S. K. & Les, C. A histological technique for detecting the cryptic preservation of erythrocytes and soft tissue in ancient human skeletonized remains. American journal of physical anthropology 152, 566–568 (2013).

49. Cattaneo, C., Gelsthorpe, K., Phillips, P. & Sokol, R. J. Blood in ancient human bone. Nature 347, 339–339 (1990).

50. Hortolà, P. SEM analysis of red blood cells in aged human bloodstains. Forensic science international 55, 139–159 (1992).

51. Loy, T. H. Prehistoric blood residues: detection on tool surfaces and identification of species of origin. Science 220, 1269–1271 (1983).

52. Cattaneo, C., Gelsthorpe, K., Phillips, P. & Sokol, R. J. Detection of blood proteins in ancient human bone using ELISA: a comparative study of the survival of IgG and albumin. International Journal of Osteoarchaeology 2, 103–107 (1992).

53. Fournier, P.-E., Drancourt, M., Aboudharam, G. & Raoult, D. Paleomicrobiology of Bartonella infections. Microbes and infection 17, 879–883 (2015).

54. Spyrou, M. A., Bos, K. I., Herbig, A. & Krause, J. Ancient pathogen genomics as an emerging tool for infectious disease research. Nature Reviews Genetics 1 (2019).

55. Barnett, R. & Larson, G. A phenol–chloroform protocol for extracting DNA from ancient samples. in Ancient DNA 13–19 (Springer, 2012).

56. Diatta, G. et al. Prevalence of Bartonella quintana in patients with fever and head lice from rural areas of Sine-Saloum, Senegal. The American journal of tropical medicine and hygiene 91, 291–293 (2014).

57. Gescher, D. M. et al. A view on Bartonella quintana endocarditis—confirming the molecular diagnosis by specific fluorescence in situ hybridization. Diagnostic microbiology and infectious disease 60, 99–103 (2008).

