## Supplementary figures for "A 2,000-year-old specimen with intraerythrocytic *Bartonella quintana*"

1. IHU Méditerranée Infection, Marseille, France.
2. Aix-Marseille-Université, IRD, MEPHI, IHU Méditerranée Infection, Marseille, France.
3. Aix-Marseille Univ, CNRS, EFS, ADES, Marseille, France.
4. Hue University of Medicine and Pharmacy, Hue, Vietnam.
5. Ville de Besançon DPH, CNRS, UMR 6298 ArTeHiS, France.
6. Laboratoire d'Hématologie, Hôpital de la Timone, APHM, Marseille, France.
7. Service d'Anatomopathologie, Assistance Publique des Hôpitaux de Marseille, Marseille, France.
8. Aix-Marseille-Université, UFR Odontology, Marseille, France.

### SUPPLEMENTARY FIGURES

Supplementary figure 1

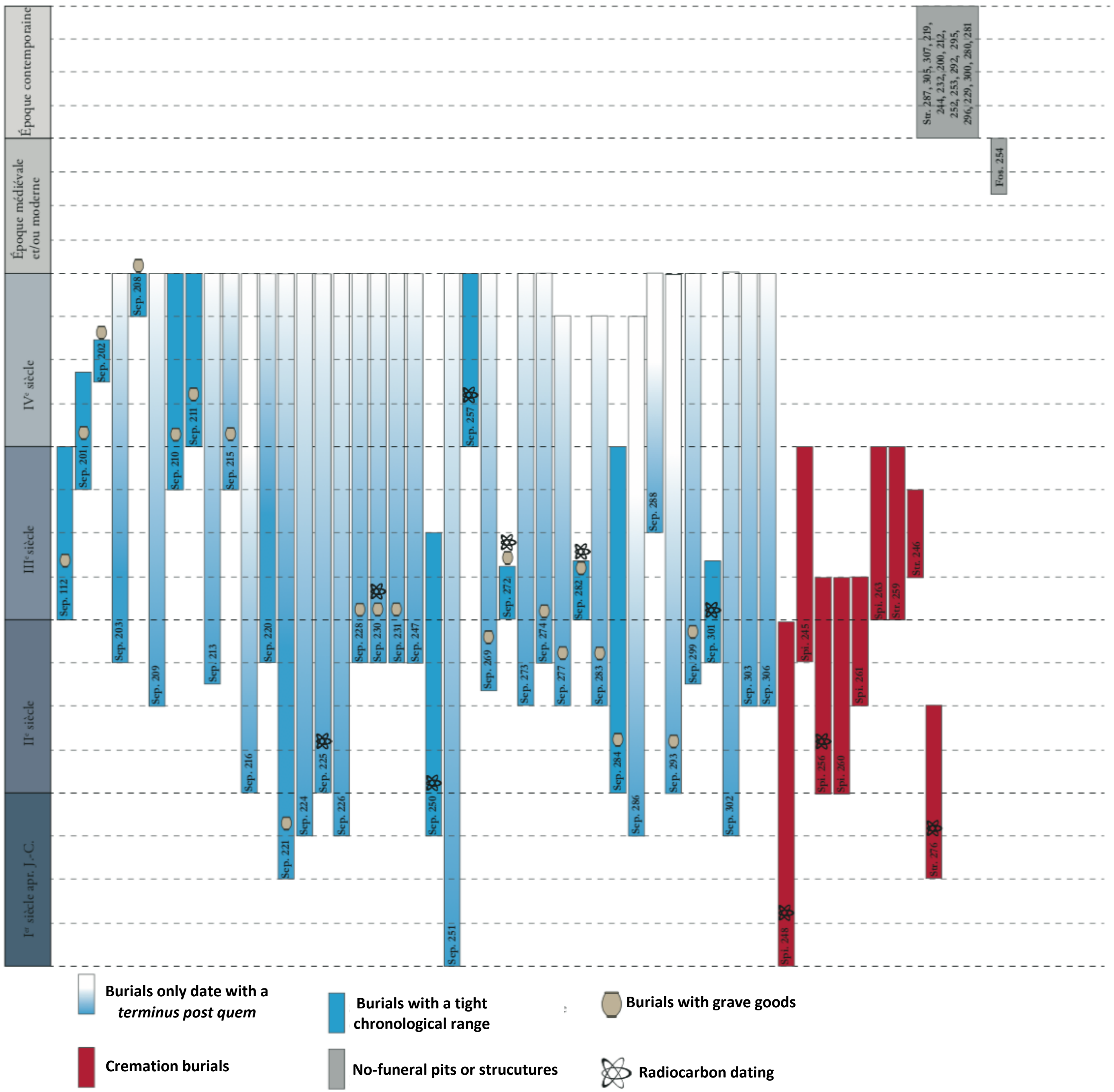

**Supplementary figure 1. synoptic view of archaeological structures including burials dating**

Supplementary figure 2

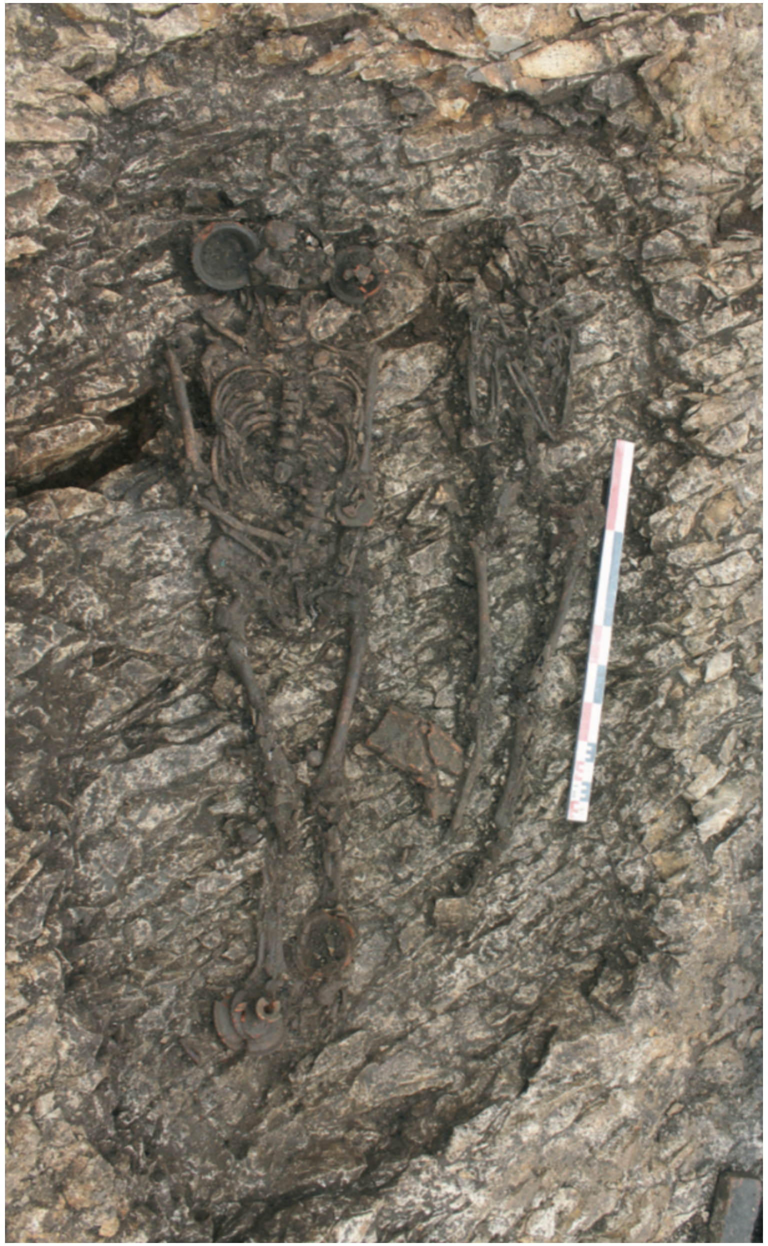

**Supplementary figure 2. General view of Sep. 230 (Ind. 21 (Left) and Ind. 20 (Right))**

Supplementary figure 3

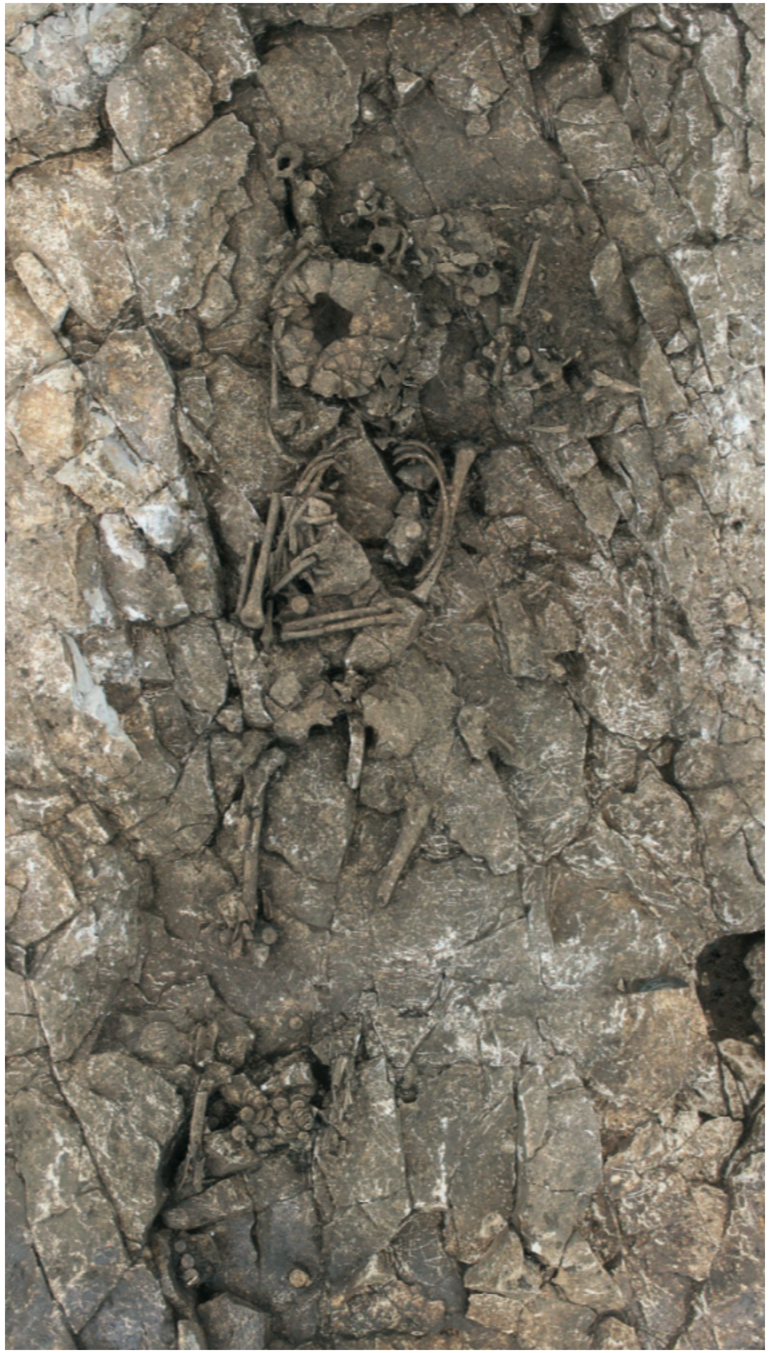

**Supplementary figure 3. General view of Sep. 283 (Ind. 33)**

Supplementary figure 4

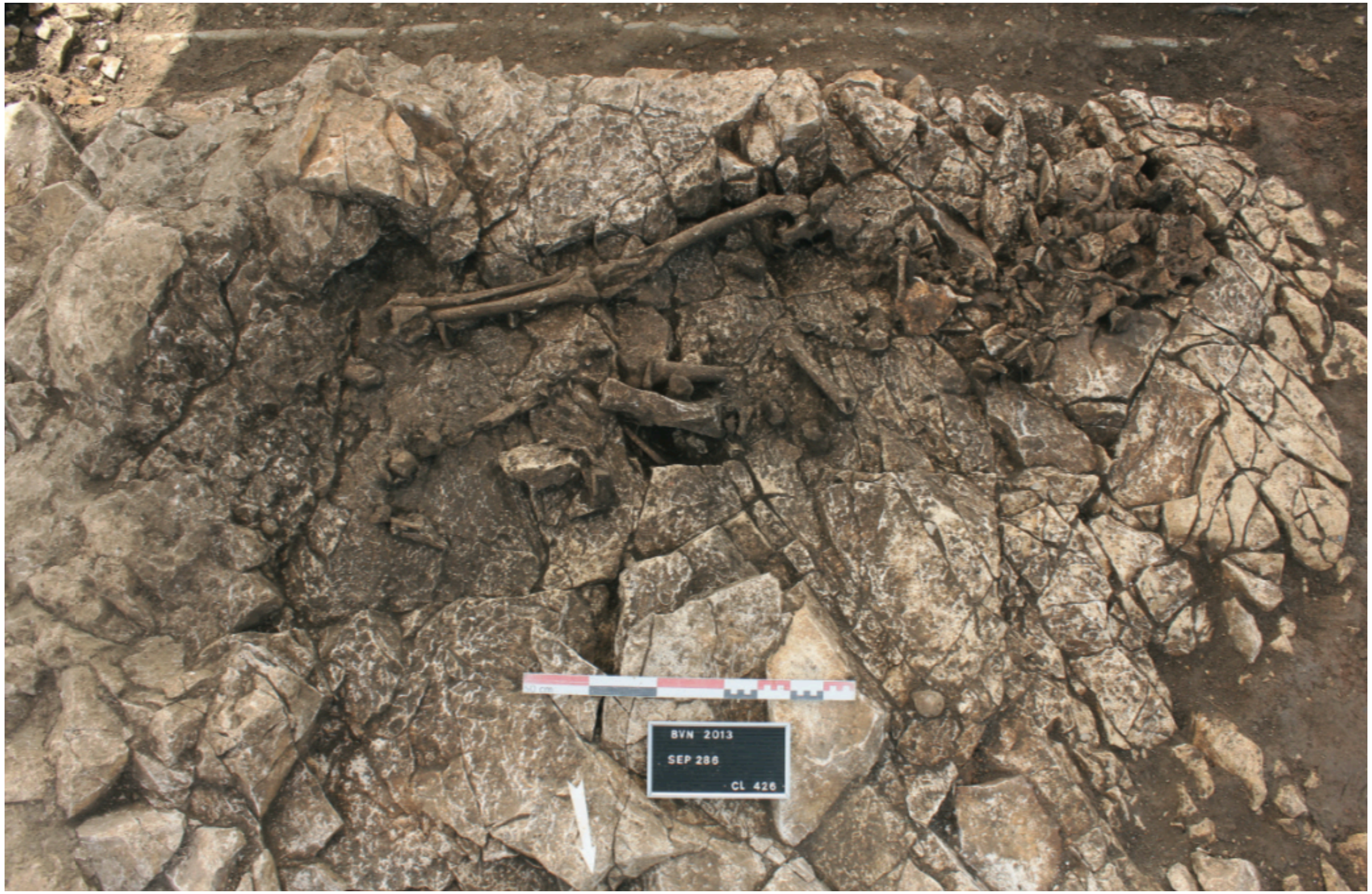

**Supplementary figure 4. General view of Sep. 286 (Ind. 35)**

Supplementary figure 5

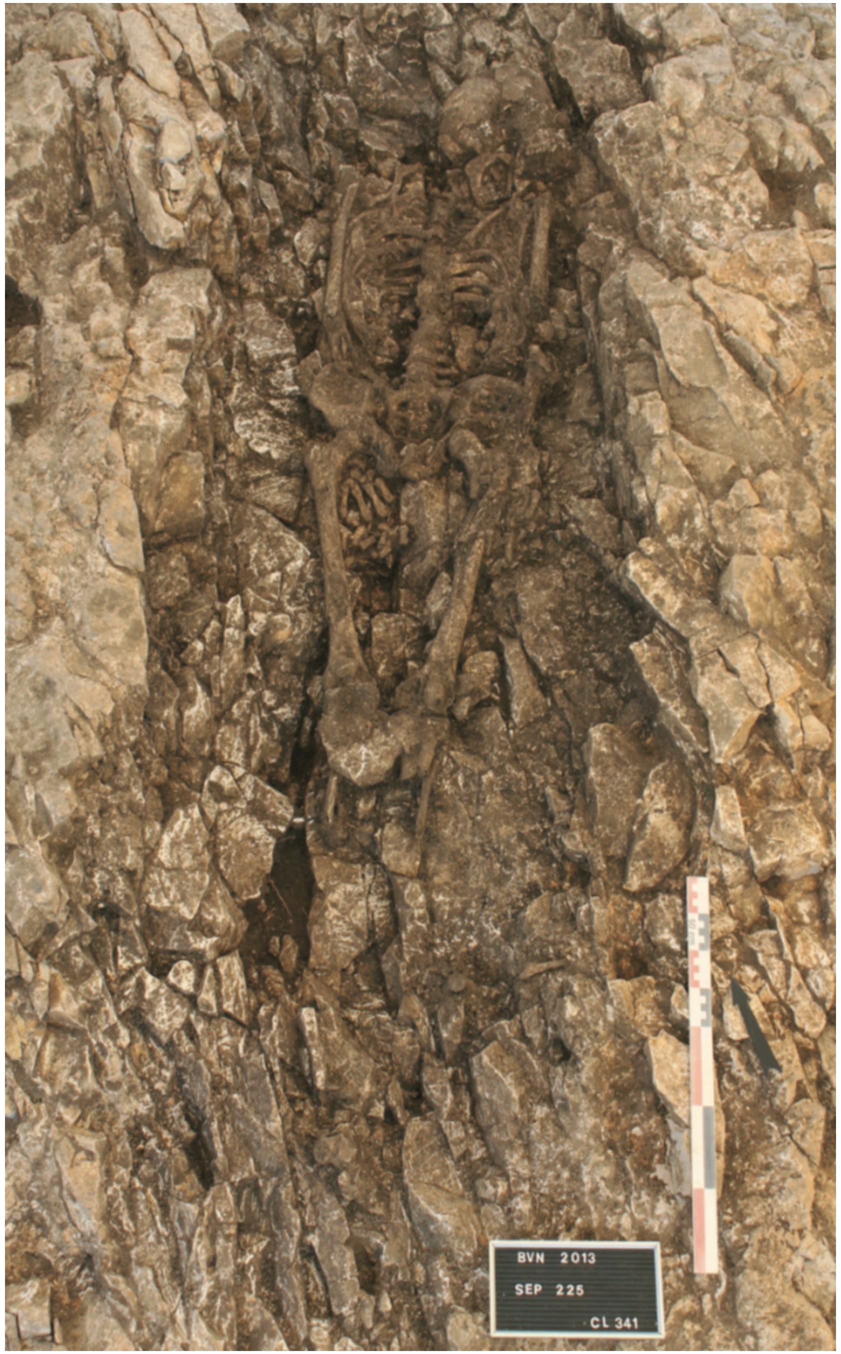

**Supplementary figure 5. General view of Sep. 225 (Ind. 17)**

**Reference : Bassi M-L, 2015. Rapport de fouilles archéologique, Besançon (Doubs), 2 rue de Vesoul, accès nord gare Viotte. L'espace funéraire Gallo-romain de la Viotte.**
